# PVTAD: ALZHEIMER’S DISEASE DIAGNOSIS USING PYRAMID VISION TRANSFORMER APPLIED TO WHITE MATTER OF T1-WEIGHTED STRUCTURAL MRI DATA

**DOI:** 10.1101/2023.11.17.567617

**Authors:** Maryam Akhavan Aghdam, Serdar Bozdag, Fahad Saeed, Alzheimer’s Disease Neuroimaging Initiative

## Abstract

Alzheimer’s disease (AD) is a neurodegenerative disorder, and timely diagnosis is crucial for early interventions. AD is known to have disruptive local and global brain neural connections that may be instrumental in understanding and extracting specific biomarkers. Previous machine-learning approaches are mostly based on convolutional neural network (CNN) and standard vision transformer (ViT) models which may not sufficiently capture the multidimensional local and global patterns that may be indicative of AD. Therefore, in this paper, we propose a novel approach called PVTAD to classify AD and cognitively normal (CN) cases using pretrained pyramid vision transformer (PVT) and white matter (WM) of T1-weighted structural MRI (sMRI) data. Our approach combines the advantages of CNN and standard ViT to extract both local and global features indicative of AD from the WM coronal middle slices. We performed experiments on subjects with T1-weighed MPRAGE sMRI scans from the ADNI dataset. Our results demonstrate that the PVTAD achieves an average accuracy of 97.7% and F1-score of 97.6%, outperforming the single and parallel CNN and standard ViT architectures based on sMRI data for AD vs. CN classification.

## 1. INTRODUCTION

World Health Organization (WHO) estimates that over 55 million people are currently living with dementia worldwide [1]. Alzheimer’s Disease (AD) is the most common type of dementia, accounting for 60-70% of cases [1]. It causes memory loss, progressive cognitive decline, and behavioral changes. While there is currently no cure for the disease, early diagnosis and prompt treatment can improve the overall quality of life and may help people with AD to live for a longer period. The clinical AD diagnosis primarily relies on cognitive, functional, and behavioral tests, which leads to AD diagnosis after symptoms have manifested, which may be too late for early interventions [2]. Therefore, biomarkers specific to AD for early and accurate AD diagnosis before behavioral symptoms are urgently needed [3], [4]. Over the past decades, there is increased interest in using imaging techniques, such as structural magnetic resonance imaging (sMRI) for diagnosis of various neuro-disorders [5], [6]. Research has demonstrated that changes in venous density within the brain’s white matter (WM) are indicators of cognitive impairment in older individuals [7]. MRI technique can measure these changes and illustrate the structural damage in the brain caused by these conditions [7]. The human brain is a complex and interconnected network of regions, and analyzing the high-dimensional WM of sMRI data are not possible with conventional means. Therefore, there is an urgent need for computer-aided diagnosis (CAD) systems that can assist in early, and precise AD diagnosis.

Machine-learning (ML) algorithms are a core component of CAD systems, enabling identification of patterns in sMRI data that may not be easily detected by naked eye. In most previous studies, researchers employed convolutional neural network (CNN) and standard vision transformer (ViT) models to extract features from sMRI data for AD diagnosis [8]–[13]. The convolutional layers in a CNN extract local features from the specific area of the input image. In contrast, standard ViT can extract global features from the whole image using attention mechanism. However, AD is known to have disruptive local and global connections [14], which may not be well captured with CNN based models alone (better for local) or may not be well captured with standard ViT models alone (which may work well for global patterns). Therefore, there is an urgent need for designing and developing ML models that can capture both local and global patterns in a synergistic manner.

The contribution of this paper are as follows:

1. In this paper, we propose a framework called PVTAD for AD vs. cognitively normal (CN) classification by applying pretrained pyramid vision transformer (PVT) [15] to extract both local and global features indicative of AD from the WM coronal middle slices of T1-weighted sMRI data.
2. We extracted all the 155 subjects that are T1-weighed MPRAGE sMRI scans available from the Alzheimer’s Disease Neuroimaging Initiative (ADNI) database. To prepare data for analysis, we extracted the WM segmentation from each 3D sMRI data and converted them into a set of 2D coronal slices. We only extracted the 30 middle coronal slices from each subject, resulting in a dataset of 4,650 2D coronal slices.
3. To showcase the power of our PVTAD model, we also implemented a parallel ResNet-18 and ViT-Tiny architecutre which can sperately extract the local and global features from the imaging data. While this parallel ResNet-18 and ViT-Tiny performed better than only ResNet-18 or only ViT-Tiny models, our prooposed PVTAD model exhibits suprior performance across all metrics.

The rest of the paper is organized as follows: In section 2, we provide the related work. In section 3, we discuss our proposed approach, which includes dataset, preprocessing steps, and our proposed approach. In section 4, we cover the experimental settings and results. Finally, in section 5, we provide our conclusions.

## 2. RELATED WORK

There is an increased interest in using ML methods for early AD diagnosis based on sMRI data. Suk et al. [8] proposed a method that combines sparse regression and CNN based on region of interests of sMRI data and achieved an accuracy of 91.02% for the AD vs. CN classification. Li and Liu [9] proposed a classification method using multiple cluster dense CNNs based on patches extracted from each local region of sMRI data, and the AD/CN classification reached an accuracy of 89.5%. Ebrahimi et al. [10] applied various 2D CNN architectures and 2D CNN-long short-term memory (LSTM) models to sagittal, coronal, and axial slices of sMRI data. Also, in this research, they proposed a voxel-based CNN method achieving an accuracy of 96.88% for AD vs. CN classification. Islam and Zhang [11] applied a 2D CNN to coronal slices of sMRI data for the AD classification problem and the proposed approach obtained an accuracy of 94.97%. Lyu et al. [12] improved the standard patch operation in vanilla (standard) ViT using a slice-wise convolution embedding method and achieved an accuracy of 96.8% for AD vs. CN classification task. Li et al. [13] proposed a novel approach integrating CNNs and transformers. While multiple ML models have been proposed, most of them have focused on either extracting local features (using CNN) or global features (using ViT) leading to models that may not be generalizable. Furthermore, most of the models are not available as open source which makes it difficult to evaluate the performance of these models.

## 3. MATERIALS AND METHODS

### 3.1 Dataset and preprocessing

#### Inclusion criteria

In this paper, we used 155 subjects (70 AD and 85 CN) who have T1-weighted magnetization prepared rapid gradient echo (MPRAGE) sMRI scans from the ADNI database (adni.loni.usc.edu) as they provide high spatial resolution and tissue contrast [16], making them ideal for studying changes in brain tissues in AD. Table 1 summarizes demographic characteristics and clinical information of selected subjects.

**Table 1.** Demographic and clinical information from the ADNI dataset.

| Data | Group | N | Age | Gender<br>[M/F] | MMSE |
| --- | --- | --- | --- | --- | --- |
| sMRI | CN | 85 | 72.13±8.4 | 50/35 | 28.4±1.24 |
|  | AD | 70 | 74±9.25 | 45/25 | 23.5±2.15 |

N: Number; M: Male; F: Female; MMSE: Mini-mental state examination; The value following ‘±’ is standard deviation.

In this study, we used statistical parameter mapping - (SPM12) (fil.ion.ucl.ac.uk/spm/software/spm12) to perform preprocessing operations on the T1-weighted sMRI scans. First, we segmented the sMRI data into gray matter, WM, and cerebrospinal fluid. Then, we applied spatial normalization and smoothing to the WM images. During the segmentation preprocess step, we set the bias regularization to very light regularization and bias cut-off FWHM to 60 mm. Also, we used the ICBM space template for affine regularization on all samples. Next, we spatially normalized the WM images to the MNI space. After this step, the shape of the data samples was 79 × 95 × 79. In addition, we considered the voxel size equal to 2 × 2 × 2 mm^3^. Finally, we used a Gaussian kernel of 8 mm FWHM to smooth the normalized WM images.

### 3.2 sMRI data decomposition from 3D to 2D

The 3D sMRI data can be composed of three slice orientations, including sagittal, coronal, and axial. Typically, coronal view provides a more comprehensive and clear view of the brain’s structures compared to the other two directions. Additionally, coronal slices can encompass three crucial tissues associated with AD, namely the cerebral cortex, the ventricle, and the hippocampus [17]. Therefore, we chose the coronal view for the selection of key slices in this study. We decomposed each 3D preprocessed sMRI data into 95 2D coronal slices using a data converter tool [18]. We extracted only the 30 middle coronal WM slices from each of the 155 subjects rather than all 95 coronal slices because 30 middle slices give a clearer view than other slices. This produced a 4,650 2D WM slices (155 subjects × 30 slices corresponding to each subject) in portable network graphic (PNG) format, including 2,550 CN slices (85 subjects × 30 slices) and 2,100 AD slices (70 subjects × 30 slices).

### 3.3 PVTAD framework

In this paper, we utilized the PVT model [15] to analyze WM coronal middle slices. PVT is an extension of ViT [19] architecture with a hierarchical (pyramid) feature extractor. It uses multiple levels of transformers [20] at different scales to capture both local and global features of WM coronal middle slices in a unified architecture. In other words, the model processes sMRI coronal middle slices as a sequence of variable-size patches, which are linearly embedded. The PVT consists of multiple stages, each extracting features at different resolution levels [15]. The Fig. 1 shows the overall PVT architecture applied to the coronal mid-slices of WM of T1-weighted sMRI data. Fig. 1 illustrates the four-stage process of PVT that produces feature maps of coronal slices at different scales. Each stage has an architecture consisting of a patch embedding layer and *L*_*i*_ transformer encoder layers. First, the model takes a sMRI coronal slice with dimensions *H*×*W*×3 and divides it into 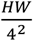 patches of size 4×4×3. Each patch is then flattened and projected linearly to create embedded patches of size 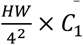. These embedded patches and position embedding are passed through a transformer encoder with *L*_1_ layers to produce a feature map *F*_1_of size 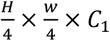. Using *F*_1_as input, this process is repeated to obtain *F*_2_, *F*_3_, and *F*_4_feature maps of coronal slices. 8, 16, and 32 are the stride of each subsequent feature maps [15].

**Fig 1.**
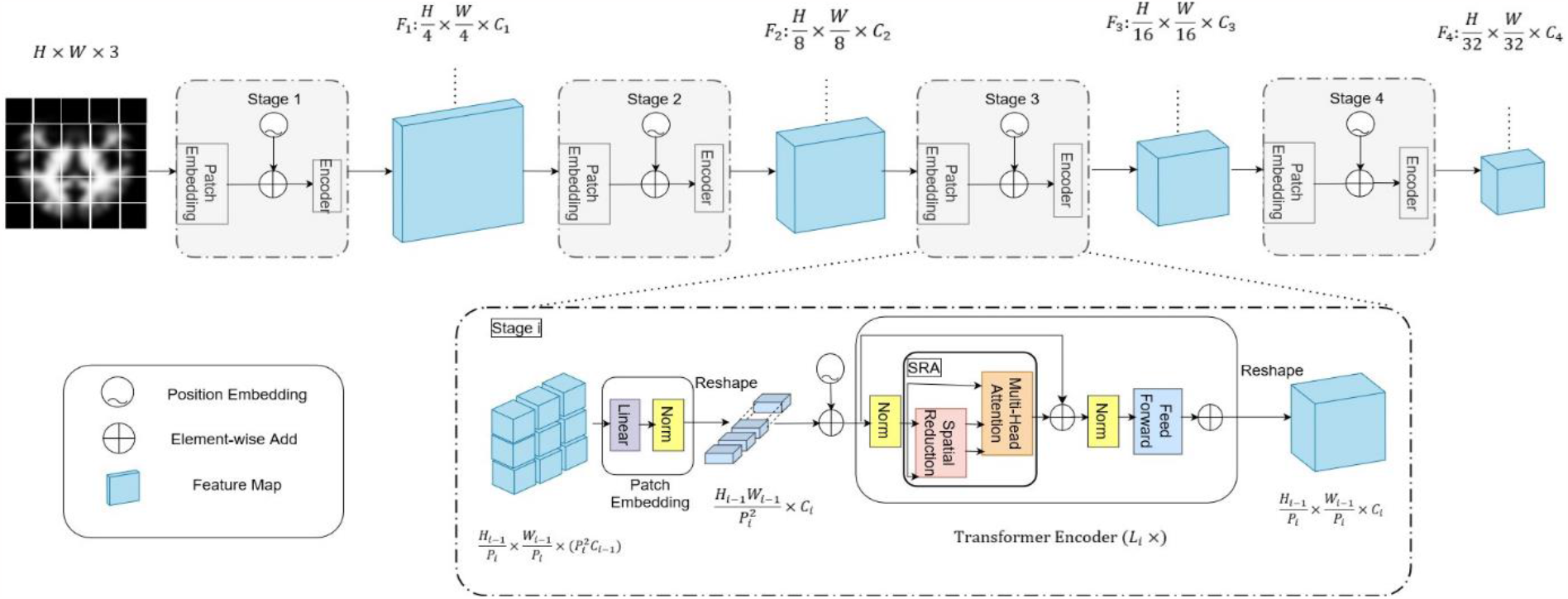
Overall architecture of PVT architecture applied to WM of T1-weighted coronal middle slices of sMRI data.

PVT employs a progressive shrinking strategy through patch embedding layers to regulate the scales of feature maps [15]. This involves equally dividing the input feature map 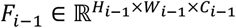 into 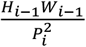 patches, where the patch size is denoted by *P*_*i*_[15]. Afterward, each patch is flattened and projected into a *C*_*i*_-dimensional embedding. The embedded patches form a shape of 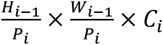, where the height and width are *P*_*i*_times smaller than the original input size [15]. To handle high-resolution feature maps, a spatial-reduction attention (SRA) layer is replaced by multi-head attention (MHA) layer [20] in stage *i* of the transformer encoder. The SRA layer takes in a query, a key, and a value (denoted as Q, K, and V, respectively) and produces a refined feature as output. The formula for SRA is expressed as equations (1) and (2) [15]:

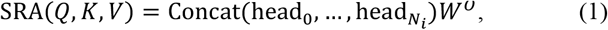

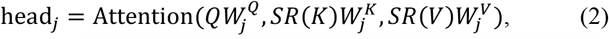

where 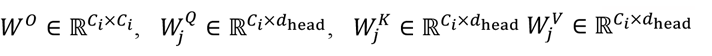, and *N*_*i*_denotes the head number of the attention layer. The dimensions of each head (*d*_head_) is 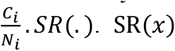 is formulated as equation (3) [15]:

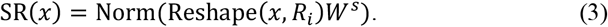

where 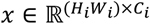 and *R*_*i*_represent an input sequence and the reduction ratio of the attention layers, respectively. Operation of Reshape(x, *R*_*i*_) reshapes x to 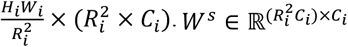 and Norm(·) are linear projection and layer normalization, respectively.

## 4. EXPERIMENTS AND EVALUATIONS

### 4.1 Experimental settings

We shuffled the 3D preprocessed WM of T1-weighted sMRI data and created five training datasets, including 70% of subjects, five validation datasets, including 10% of subjects, and five test datasets, including 20% of subjects for AD and CN groups at the subject level. This ensured no overlap among the samples in training, validation, and test datasets. Then, as we mentioned in section 3.2, we converted each 3D WM of T1-weighted sMRI data to a set of 2D coronal slices using a data converter tool [18], and we extracted only the 30 middle coronal WM slices from each of the 155 subjects. This dataset contained 4,650 2D coronal slices (2,100 AD and 2,550 CN slices).

In this study, we used PVT-Tiny model pretrained on ImageNet [21] at a resolution of 224×224 [22]. Moreover, we added a fully connected classification layer to PVT architecture for our binary classification task (AD vs. CN). To fine-tune the pretrained PVT-Tiny model, we considered freezing all the weights except for those in the final classification layer. During the fine-tuning process, we adjusted the model for 20 epochs and initialized for adaptive moment estimation (Adam) optimizer [23] with a learning rate of 0.001. We implemented the code using Keras [24] and TensorFlow Image Models (tfimm) library [25]. In addition, we conducted our experiments on a machine with an Intel(R) Xeon(R) Gold 6152 CPU @ 2.10GHz with 125GB RAM. The GPU used is NVIDIA TITAN Xp.

### 4.2 Experimental results

Fig. 2 (a) and Fig. 2 (b) show training and validation accuracy and loss for PVTAD framework on WM coronal mid-slices, respectively. In addition, to demonstrate the effectiveness of the PVTAD framework, we implemented a parallel ResNet-18 [26], a CNN-based model, and ViT-Tiny [19], a standard ViT-based model, as a basis for comparison. Fig. 2 (c) displays the receiver operating characteristic (ROC) curve of the PVTAD model and the single and parallel ResNet-18 and ViT-Tiny architectures. This figure shows PVTAD model achieves an area under the curve (AUC) of 98%, outperforming the single and parallel ResNet-18 and ViT-Tiny architectures. Table 2 presents the experimental results of this paper based on WM coronal mid-slices. According to this table, the PVTAD framework achieves an average accuracy of 97.7%, sensitivity of 97.15%, specificity of 98.16%, precision of 98.02, and F1-score of 97.6%, outperforming previous studies [10], [12]. Moreover, while parallel ResNet-18 and ViT-Tiny performed better than only ResNet or only ViT models, our proposed PVTAD model exhibits superior performance across all metrics.

**Table 2.** Experimental results for PVTAD framework in comparison with single and parallel ResNet-18 and ViT-Tiny models for AD vs. CN classification on ADNI dataset based on five-fold cross validation.

| Study | Modality | Model | ACC | SEN | SPE | PRE | F1 | AUC |
| --- | --- | --- | --- | --- | --- | --- | --- | --- |
| [10] | sMRI (Coronal View) | ResNet-18 | 81.25 | - | - | - | - | - |
| [12] | sMRI (Coronal View) | ResNet-18 | 88.5 | 100 | - | 75.9 | 86.3 | - |
|  |  | ViT-Tiny | 95.3 | 94.4 | - | 90 | 93.2 | - |
| This paper | sMRI (WM Coronal View) | ResNet-18 | 71.06 | 62.91 | 78.7 | 73.7 | 67.61 | 72 |
|  |  | ViT-Tiny | 96.23 | 95.91 | 96.53 | 96.3 | 96.1 | 95 |
|  |  | Parallel ResNet-18 and ViT-Tiny | 96.87 | 96.62 | 97.19 | 96.92 | 96.76 | 96 |
|  |  | **PVTAD model** | **97.7** | **97.15** | **98.16** | **98.02** | **97.6** | **98** |

ACC: Accuracy; SEN: Sensitivity; SPE: Specificity; PRE: Precision; F1: F1-score; AUC: Area under the curve.

**Fig 2.**
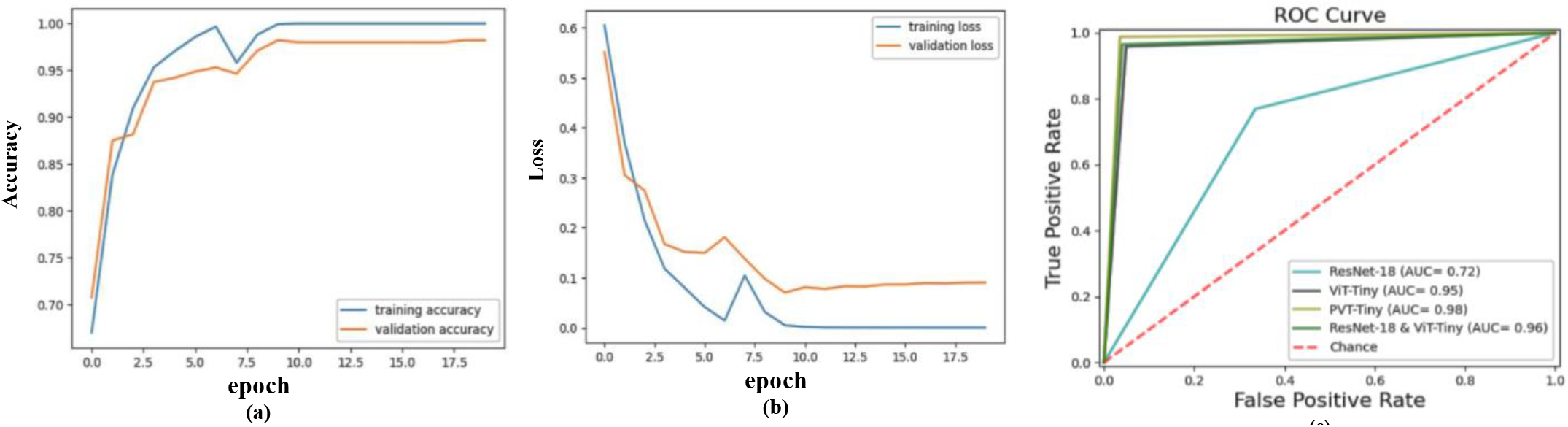
**(a)**. Training and validation accuracy for PVTAD model; **(b)** Training and validation Loss for PVTAD model; **(c)** ROC Curve for PVTAD and single and parallel ResNet-18 & ViT-Tiny models

## 5. CONCLUSIONS

AD is a progressive brain disorder, and timely diagnosis helps clinician develop patient-specific treatment plans that can address the symptoms of the disease and delay cognitive decline. AD has complex disruptions in local and global neural connections in the brain which may not be fully captured by conventional ML models such as CNN and ViT. In this paper, to better capture these complex patterns in AD and improve diagnosis performance, we designed and developed a framework called PVTAD using pretrained PVT model. Our proposed framework can capture both local and global patterns indicative of AD from WM coronal middle slices of T1-weighted sMRI data. Our experiments demonstrate that the proposed architecture leads to improve performance in AD vs. CN classification task. Moreover, this contrasts with parallel CNN and standard ViT model which can extract local and global features separately, but fail to integrate them effectively, resulting in suboptimal feature representation. To evaluate the model, we performed experiments on 155 subjects who have T1-weighted MPRAGE sMRI from the ADNI dataset. PVTAD framework achieves an average accuracy of 97.7%, sensitivity of 97.15%, specificity of 98.16%, precision of 98.02%, and F1-score of 97.6%, outperforming the single and parallel CNN and standard ViT architectures.

## 6. COMPLIANCE WITH ETHICAL STANDARDS

This research study was conducted retrospectively using human subject data made available in open access by ADNI dataset. Ethical approval was not required and approval documents for can be accessed via (adni.loni.usc.edu).

## 7. ACKNOWLEGMENTS

This material is based upon work supported by the National Institutes of Health (NIH) grant number R01GM134384 and partly by National Science Foundation (NSF) grant NSF TI-2213951. The content is solely the responsibility of the authors and does not necessarily represent the official views of the National Institutes of Health (NIH) or the National Science Foundation (NSF).

